# Identification of biological processes and signaling pathways in the stretched nucleus pulposus cells

**DOI:** 10.1101/2021.11.23.469730

**Authors:** Min Zhang, Jing Wang, Hanming Gu

**Affiliations:** SHU-UTS SILC School, Shanghai University, Shanghai, China

**Author notes:** Corresponding author: Hanming Gu, SHU-UTS SILC School, Shanghai University, Shanghai, China.

## Abstract

Low back pain is mostly caused by disc degeneration, which is due to the alterations in the osmotic pressure of nucleus pulposus cells. However, the knowledge about the mechanism and therapies for disc degeneration is not fully understood. Here, our objective is to identify significantly changed genes and biological processes in the stretched nucleus pulposus cells. The GSE175710 dataset was originally produced by using the Illumina HiSeq 4000 (Rattus norvegicus). The KEGG and GO analyses indicated that “MAPK signaling”, “TNF signaling”, “IL17 signaling”, and the “NF-κB signaling pathway” are mostly affected in the stretched nucleus pulposus cells. Moreover, we identified several genes according to the PPI network such as Mmp9, Cxcl12, Col1a1, and Col3a1 in the stretched nucleus pulposus cells. Thus, our study provides further insights into the study of disc deterioration.

## Introduction

Low back pain affects nearly 80% of the population worldwide, and most cases can be defined as intervertebral disc degeneration^1^. About two-thirds of adults suffer from back pain, and most pain is due to disc diseases^2^. Unfortunately, the current medical strategies have no significant improvement in patients such as health status and disability^3^. Furthermore, the back pain surgery for degenerative discs was found to have negatively affected the mechanics of surrounding discs^4^. The intervertebral disc separates the vertebral bodies in the spine that are strained by various factors including compression, bending, and shear^5^. Certain spinal stress is associated with disc injury, degeneration, and damage^6^. Spinal loading on the disc is also affected by various conditions including oxygen tension, pH, water content, and permeability^7^.

The lack of nucleus pulpous cells in the disc is the early sign of disc degeneration^4^. The nucleus pulpous cells maintain the extracellular matrix through secreting proteoglycans, aggrecan, and collagens^8^. A mechanical loading system that applies cyclic stretch stress can qualify the cells’ deformation and strain^9^. Since the capacity of nucleus pulposus cells is the reason for disc degeneration, it is a critical approach to restore or regenerate the nucleus pulposus cells^10^. A number of studies tested stem cell-based therapy for inhibiting disc degeneration by transplantation of mesenchymal stem cells into the vertebral disc, which has been proven to decrease disc degeneration in animal models^11^. Though disc degeneration is caused by mechanical stress on the nucleus pulposus, the mechanism and functions of nucleus pulposus cells in vitro have not been clarified^12^.

In this study, we evaluated the effects of loading stress on the nucleus pulposus cells by analyzing the RNA sequence data. We identified a bunch of DEGs and several significant biological pathways. We also introduced the gene function enrichment and created the protein-protein interaction (PPI) network and Reactome map for finding the potential interacting proteins and pathways. These functional genes and biological processes will shed light on the treatment of low back pain.

## Methods

### Data resources

The data was created by using the Illumina HiSeq 4000 (Rattus norvegicus) (Beijing TongRen Hospital, Capital Medical University, No. 1, Dongjiaomin Lane, Dongcheng District, Beijing, China). The analyzed dataset includes three groups of control cells and three groups of stretched cells.

### Data acquisition and preprocessing

We processed the raw data by the R package^13, 14^. We performed a classical t-test to identify DEGs with P< 0.001 and fold change ≥ 1 as being statistically significant.

### The Kyoto Encyclopedia of Genes and Genomes (KEGG) and Gene Ontology (GO) analyses

We performed the KEGG and GO analyses in this study by using the Database for Annotation, Visualization, and Integrated Discovery (DAVID) (http://david.ncifcrf.gov/). We set the P<.05 and gene counts >10 as the statistically significant cutoff.

### Protein-protein interaction (PPI) networks

The PPI networks were constructed and verified by using the Molecular Complex Detection (MCODE)^15^. The significant modules were created from constructed PPI networks. The biological pathways analysis was performed by Reactome (https://reactome.org/), and P<0.05 was used as the cutoff criterion.

## Results

### Identification of DEGs between the control and elongated nucleus pulposus cells

To determine the molecular mechanisms of excessive mechanical strain on nucleus pulposus cells, we analyzed the DEGs from the control and elongated nucleus pulposus cells. A total of 484 genes were identified with the threshold of P<0.05. The top ten of up-and-down-regulated genes for the control and elongated nucleus pulposus cells are shown by the heatmap and volcano plot (Figure 1). The top ten DEGs were listed in Table 1.

**Table 1.**

| **Entrez gene** | **Gene Symbol** | **Fold-change** | **Regulation** |
| --- | --- | --- | --- |
| Top 10 down-regulated DEGs |  |  |  |
| 681355 | LOC681355 | -4.308594983 | Down |
| 1240 | Cmklr1 | -4.188048629 | Down |
| 2487 | Frzb | -3.826703132 | Down |
| 103689971 | LOC103689971 | -3.823545803 | Down |
| 80168 | Mogat2 | -3.58056859 | Down |
| 2619 | Gas1 | -3.374662642 | Down |
| 7060 | Thbs4 | -3.239004297 | Down |
| 55220 | Klhdc8a | -3.198784255 | Down |
| 2637 | Gbx2 | -3.191774195 | Down |
| 25992 | Sned1 | -3.184824365 | Down |
| Top 10 up-regulated DEGs |  |  |  |
| 6348 | Ccl3 | 7.38862333 | up |
| 6352 | Ccl5 | 6.817080418 | up |
| 969 | Cd69 | 6.331380669 | up |
| 4318 | Mmp9 | 6.218007819 | up |
| 1437 | Csf2 | 6.203278857 | up |
| 445 | Ass1 | 6.153898019 | up |
| 3488 | Igfbp5 | 6.150743666 | up |
| 1440 | Csf3 | 5.895138107 | up |
| 5055 | Serpinb2 | 5.815539174 | up |
| 4319 | Mmp10 | 5.64306114 | up |

**Figure 1.**
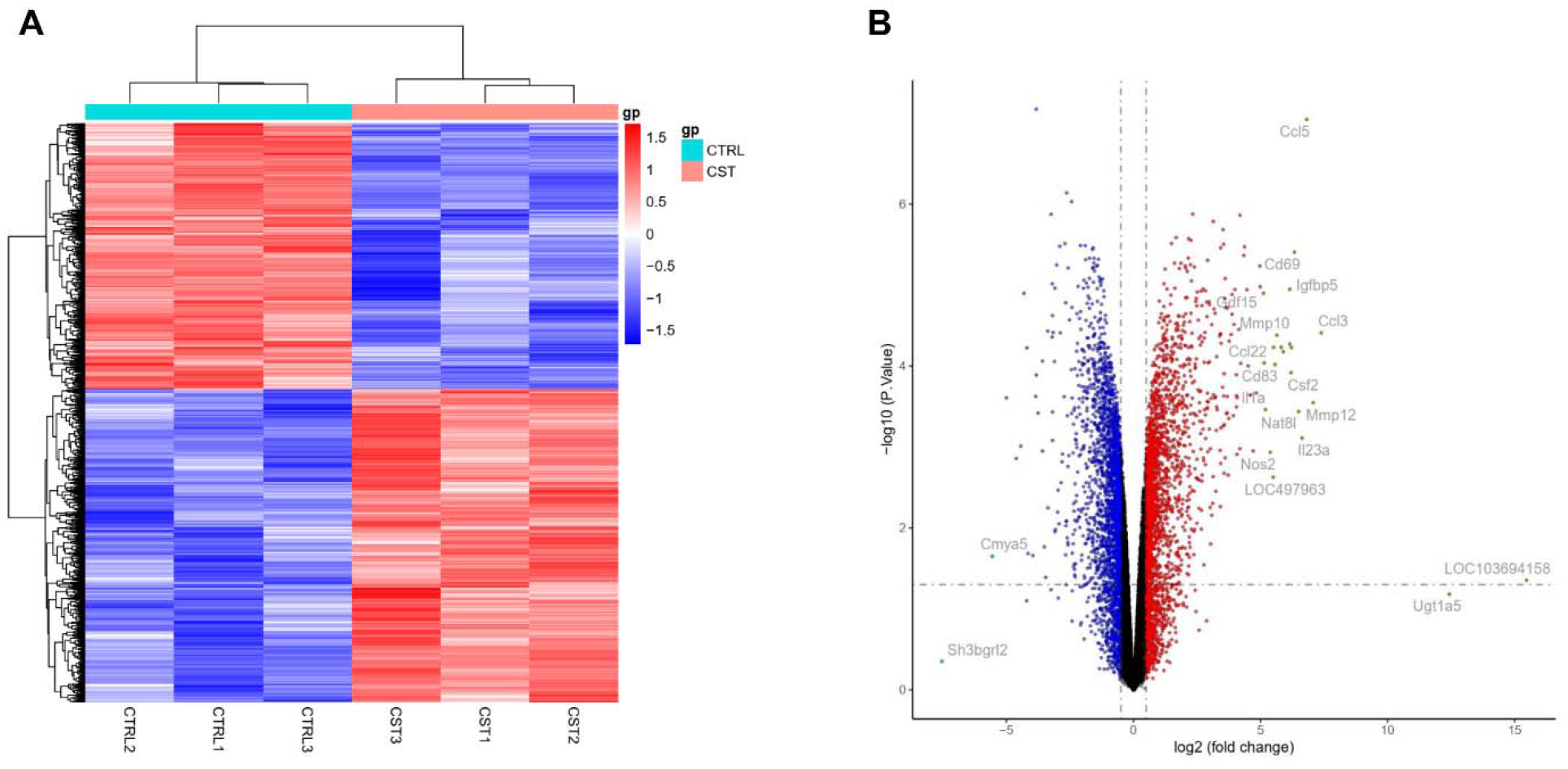
Heatmap and volcano plot between the control and elongated nucleus pulposus cells. (A) Heatmap of significant DEGs. Significant DEGs (P < 0.01) were used to create the heatmap. (B) Volcano plot for DEGs in the control and elongated nucleus pulposus cells. The mostly changed genes are highlighted by grey dots and symbols marked.

### Enrichment analysis of DEGs between the control and elongated nucleus pulposus cells

To determine the biological roles of DEGs under the conditions of excessive mechanical strain for nucleus pulposus cells, we performed the KEGG and GO analyses as previously described^16, 17^ (Figure 2). The top KEGG pathways contain “MAPK signaling pathway”, “Focal adhesion”, “TNF signaling pathway”, “Fluid shear stress and atherosclerosis”, “IL-17 signaling pathway”, “AGE-RAGE signaling pathway in diabetic complications”, “NF-kappa B signaling pathway”, “ECM-receptor interaction”, “Rheumatoid arthritis”, and “Steroid biosynthesis”. We then identified the biological processes of GO, including “Positive regulation of cell adhesion”, “Ameboidal-type cell migration”, “Extracellular matrix organization”, “Extracellular structure organization”, “External encapsulating structure organization”, “Cell-substrate adhesion”, “Leukocyte migration”, “ERK1 and ERK2 cascade”, “Regulation of ERK1 and ERK2 cascade”, and “Peptidyl-tyrosine modification”. We also identified the top ten cellular components of GO, including “Collagen-containing extracellular matrix”, “Apical part of cell”, “Lytic vacuole”, “Lysosome”, “Neuron to neuron synapse”, “Cell leading edge”, “Receptor complex”, “Asymmetric synapse”, “Postsynaptic density”, and “Early endosome”. We further identified the molecular functions of GO, including “Nucleoside-triphosphatase regulator activity”, “DNA-binding transcription activator activity, RNA polymerase II-specific”, “GTPase regulator activity”, “Phosphoric ester hydrolase activity”, “Cell adhesion molecule binding”, “Sulfur compound binding”, “Glycosaminoglycan binding”, “Heparin binding”, “Extracellular matrix structural constituent”, and “Integrin binding”.

**Figure 2.**
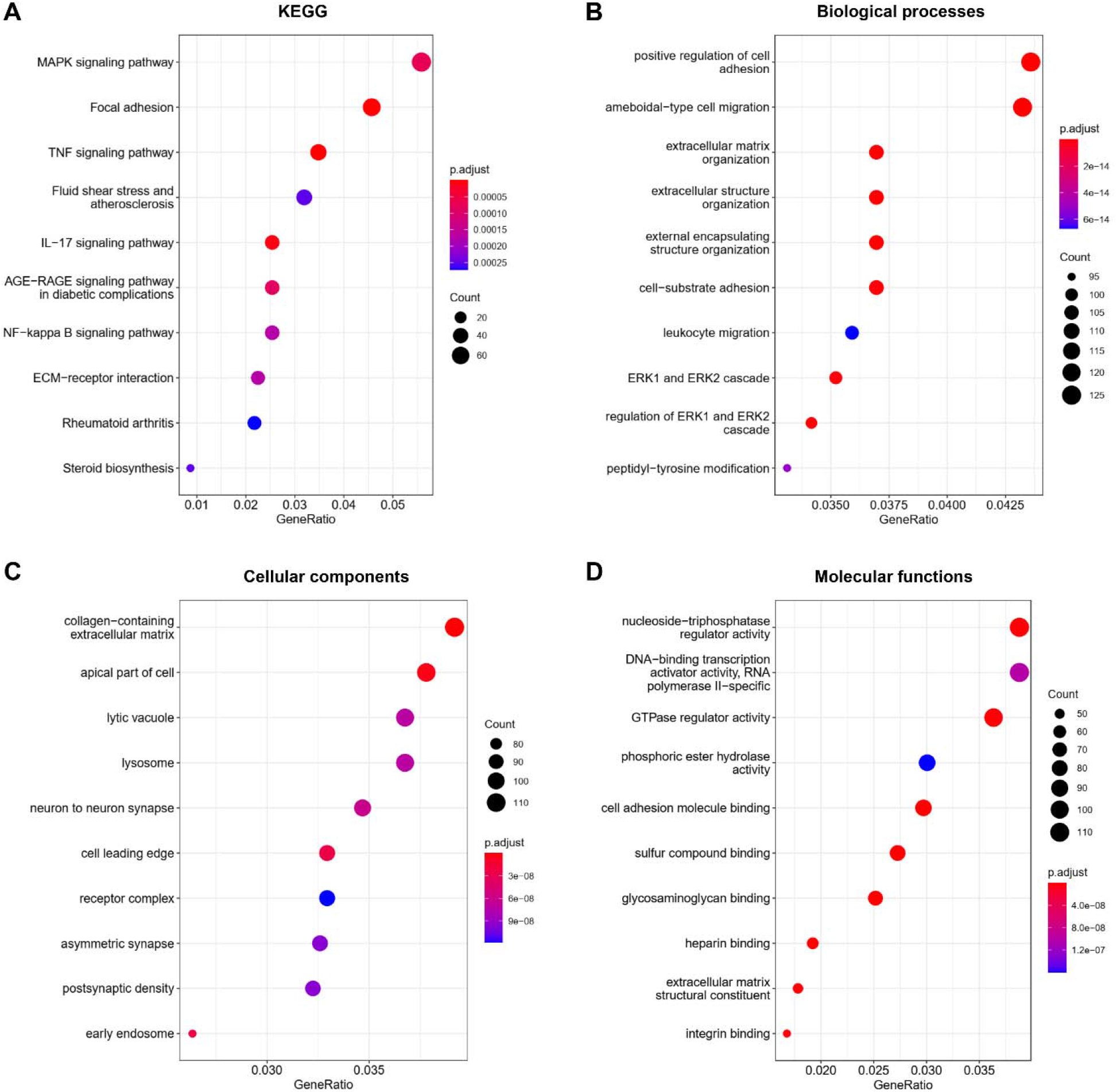
KEGG and GO analyses of DEGs. (A) KEGG analysis, (B) Biological processes, (C) Cellular components, (D) Molecular functions.

### PPI network analysis

To further determine the relationship of the DEGs, we created the PPI network by using the Cytoscape software. The criterion of combined score > 0.2 was defined to create the PPI by using the 381 nodes and 1182 edges. Table 2 showed the top ten genes with the highest degree scores. The top two modules depicted the functional annotation (Figure 3). We further analyzed the DEGs and created the biological map by Reactome (Figure 4). We identified the top ten significant pathways including “Activation of gene expression by SREBF (SREBP)”, “Regulation of cholesterol biosynthesis by SREBP (SREBF)”, “Interleukin-10 signaling”, “Collagen degradation”, “Cholesterol biosynthesis”, “Degradation of the extracellular matrix”, “Assembly of collagen fibrils and other multimeric structures”, “Metabolism of steroids”, “Extracellular matrix organization”, and “Collagen formation”.

**Table 2.**
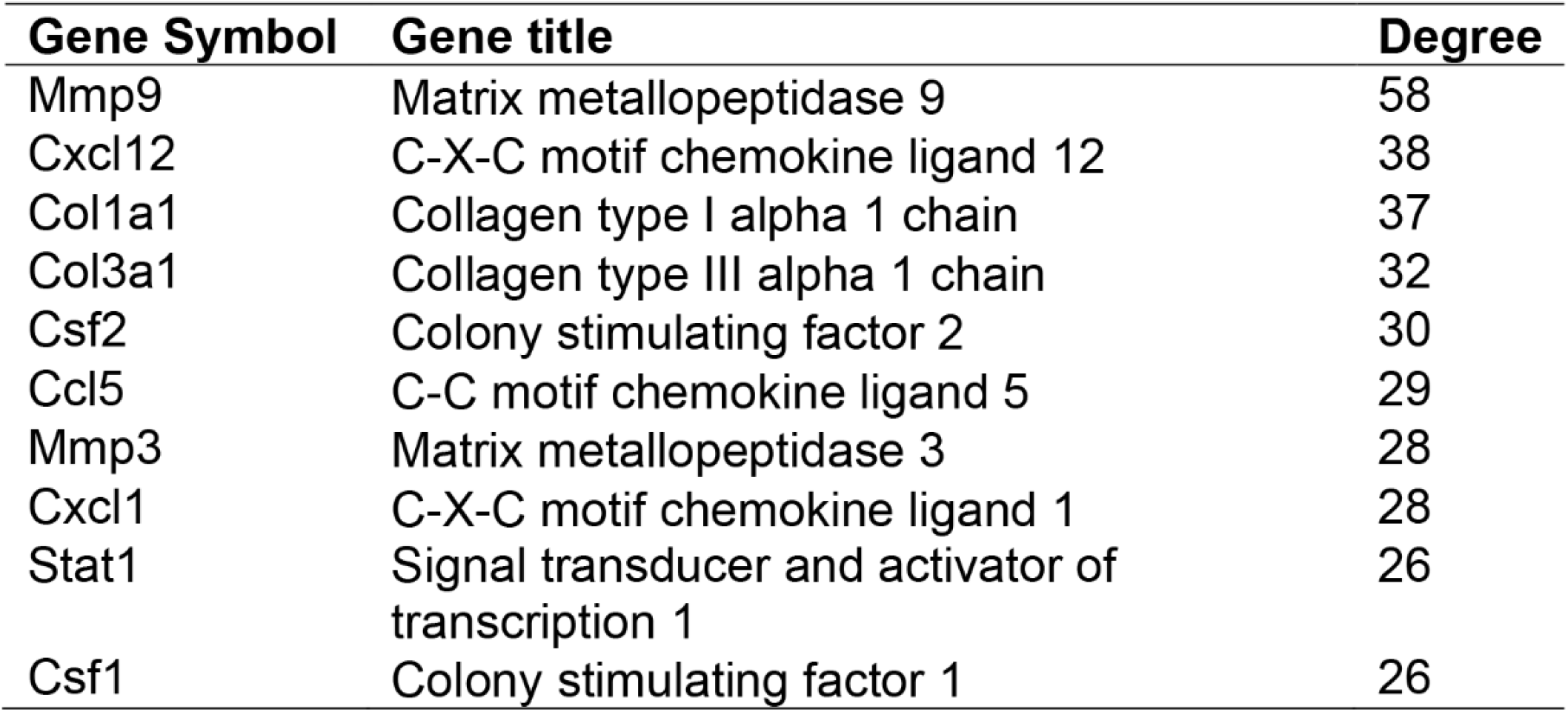
Top ten genes demonstrated by connectivity degree in the PPI network.

**Figure 3.**
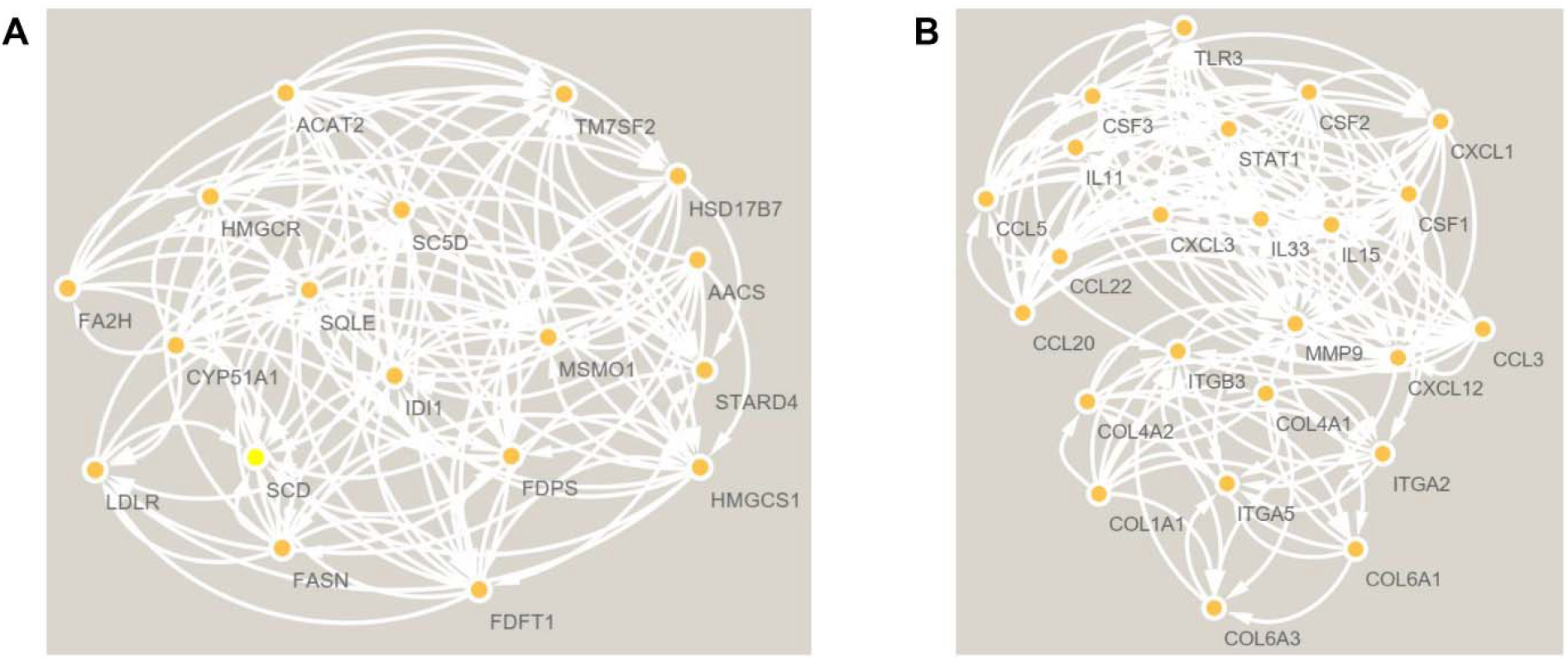
The PPI network analysis of DEGs between the control and elongated nucleus pulposus cells. Cluster 1 (A) and cluster 2 (B) were the top two clusters and were constructed by String and MCODE in Cytoscape.

**Figure 4.**
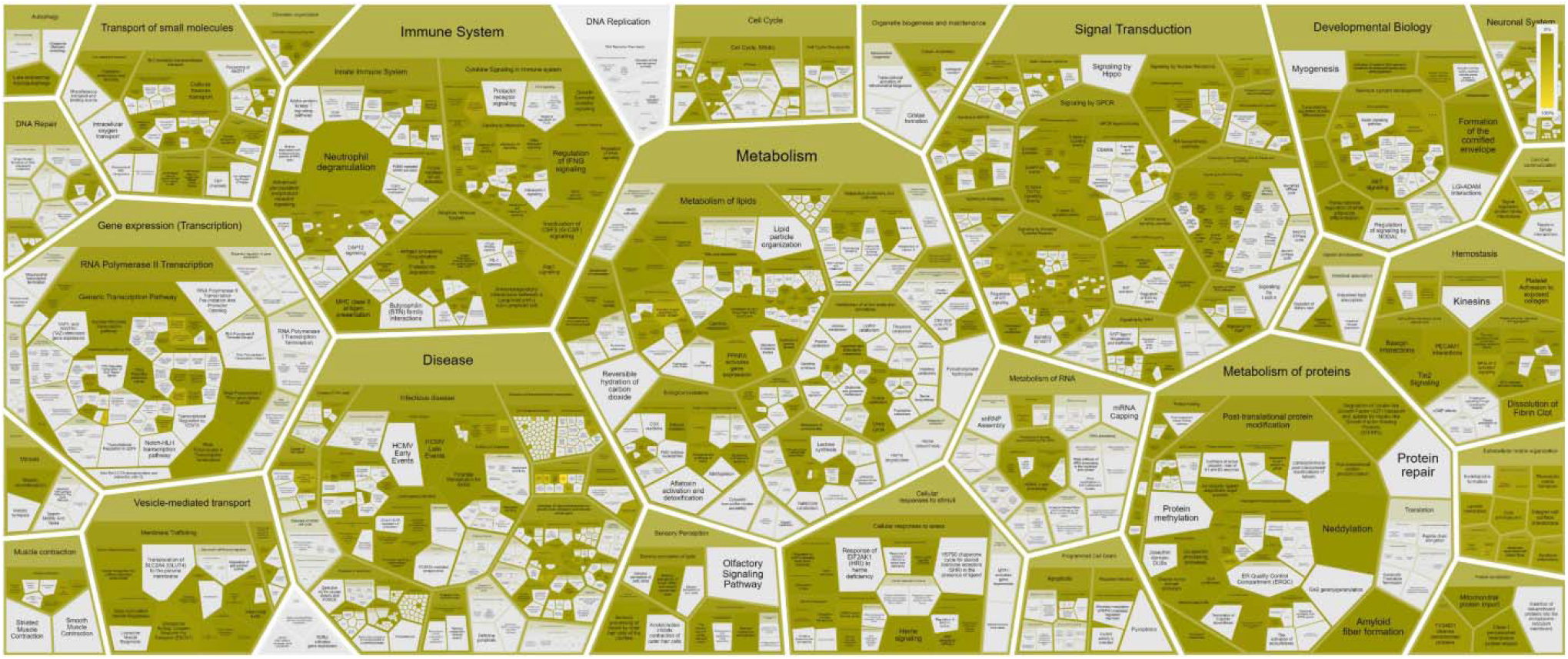
Reactome map representation of the significant biological processes between the control and elongated nucleus pulposus cells.

## Discussion

Nucleus pulposus cells are located in the ecological niches without blood vessels and nerves where the osmotic pressure is higher than the plasma and is able to hold more proteoglycans^18^. The osmotic pressure alters with the age and degeneration of the interverbal disc, which has a close relationship with the low back pain^19^.

The nucleus pulposus cells under the stress change the immune status in our studies. We found most of the immune pathways such as the MAPK signaling pathway, TNF signaling pathway, IL17 signaling pathway, and the NF-κB signaling pathway were related to the nucleus pulposus cells change. Similarly, the MAPK signaling pathway shows a critical role in controlling cell proliferation, differentiation, and apoptosis under the pathophysiological conditions of intervertebral disc^20^. At the beginning of low back pain, there are significantly higher levels of TNF in the patient’s group in comparison with the healthy group^21^. Shamji MF et al reported that the increased IL17 is closely related to the chronification of pain and the degenerated lumbar intervertebral discs^22^. NF-κB is a key inflammatory response mediator that regulates inflammation in chondrocytes and macrophages^23-25^, which can lead to osteoarthritis and rheumatoid arthritis^26, 27^. Moreover, NF-κB is able to regulate several inflammatory enzymes such as COX-2 to affect the pain sensitivity in arthritis^28, 29^. Aisha S. Ahmed et al found that NF- κB is also involved in the pain-Related neuropeptide expression in degenerative disc^30^. Interestingly, we found the nucleus pulposus cells under the stress could affect rheumatoid arthritis and steroid biosynthesis by KEGG analysis, suggesting that nucleus pulposus cells may involve in the synthesis and secretion of proteins and liposomes under the loading condition.

By analyzing the GO, we found that cell migration and cell adhesion are the most important processes in the stretched nucleus pulposus cells. C.L. Gilchrist et al found that nucleus pulposus cells are originated from the embryonic notochord that exhibits strong cell-cell interaction and adherence^31^. Weiheng Wang et al showed the allogenic nucleus pulposus cells have the anti-apoptotic and migratory ability that lead to the disc regeneration^32^. GPCR signaling and RGS proteins regulate numerous physiological and pathological processes including inflammation, bone metabolism, and pain^28, 33-36^. Strikingly, we found that nucleus pulposus cells can affect the GTPase regulator activity by analyzing the cellular components of GO, suggesting that RGS proteins may also regulate the functions of nucleus pulposus cells under the stress.

Besides the KEGG and GO analyses, we also identified the significant PPI networks and genes in nucleus pulposus cells with the loading stress. P-B Li et al found that the up-regulation of MMP9 is strongly related to the degenerated lumbar disc tissues^37^. Yang Yu et al found the CXCL12 was increased in the dorsal root ganglion after chronic compression^38^. The study by C Tilkeridis et al showed the sequence changes of the COL1A1 gene indicate a strong relationship with lumbar disc diseases^39^. Zongde et al found the decreased COL3A is associated with disc degeneration by bioinformatic methods^40^. Louise S.C. Nicol found the inhibition of GM-CSF is analgesic in neuropathic pain^41^. The circadian clocks and their downstream molecules play central roles in various physiological functions^42^ such as metabolism, immune, apoptosis, and ER stress, protein synthesis, and its disruption has been related to many diseases^43-51^. As a circadian gene-controlled gene, CCL5 regulates the most neuroprotective effects^52, 53^.

The Study by Jiangtao Wan et al showed that MMP3 is a destabilizing factor in bone diseases^54^. Rangel L Silva et al found that CXCL1 enhances nociceptor and central sensitization through CXCR2, which is a target for analgesic drugs^55^. Zhenpeng Song et al found that STAT1 leads to cancer pain as a downstream regulator of ERK signaling^56^. Zhonghui Guan et al showed the microglial expression genes that are related to pain through the CSF1 singling for survival^57^.

In conclusion, this study provided information on nucleus pulposus cells under loading conditions. MAPK signaling, TNF signaling, IL17 signaling, and NF-κB signaling are the major biological pathways in the stretched nucleus pulposus cells. This study may shed light on the treatment of low back pain and disc deterioration.

## Author Contributions

Min Zhang, Jing Wang: Methodology and Writing. Hanming Gu: Conceptualization, Methodology, Writing-Reviewing and Editing.

## Funding

This work was not supported by any funding.

## Declarations of interest

There is no conflict of interest to declare.

## Notes

### Competing Interest Statement

The authors have declared no competing interest.

